# Designing robust N-of-1 studies for precision medicine

**DOI:** 10.1101/380980

**Authors:** Bethany Percha, Edward B. Baskerville, Matthew Johnson, Joel Dudley, Noah Zimmerman

## Abstract

Recent advances in molecular biology, sensors, and digital medicine have led to an explosion of products and services for high-resolution monitoring of individual health. The N-of-1 study has emerged as an important methodological tool for harnessing these new data sources, enabling researchers to compare the effectiveness of health interventions at the level of a single individual. We have developed a stochastic time series model that simulates an N-of-1 study, facilitating rapid optimization of N-of-1 study designs and increasing the likelihood of study success while minimizing participant burden. Using simulation, we demonstrate how the number of treatment blocks, ordering of treatments within blocks, duration of each treatment, and sampling frequency affect our ability to detect true differences in treatment efficacy. We provide a set of recommendations for study designs based on treatment, outcome, and instrument parameters, and provide our simulation software as a supplement to the paper.

## Introduction

N-of-1 studies have shown great promise as a tool for investigating the effects of drugs, supplements, behavioral changes, and other health interventions on individual patients [1–7]. An N-of-1 study (Figure 1) is a multiple-crossover comparative effectiveness study of a single patient. Competing treatments are administered in blocks, within which treatment order is randomized^1^. The outcome of interest is compared across different treatment periods to find the treatment with the greatest efficacy for that specific patient.

**Figure 1:**
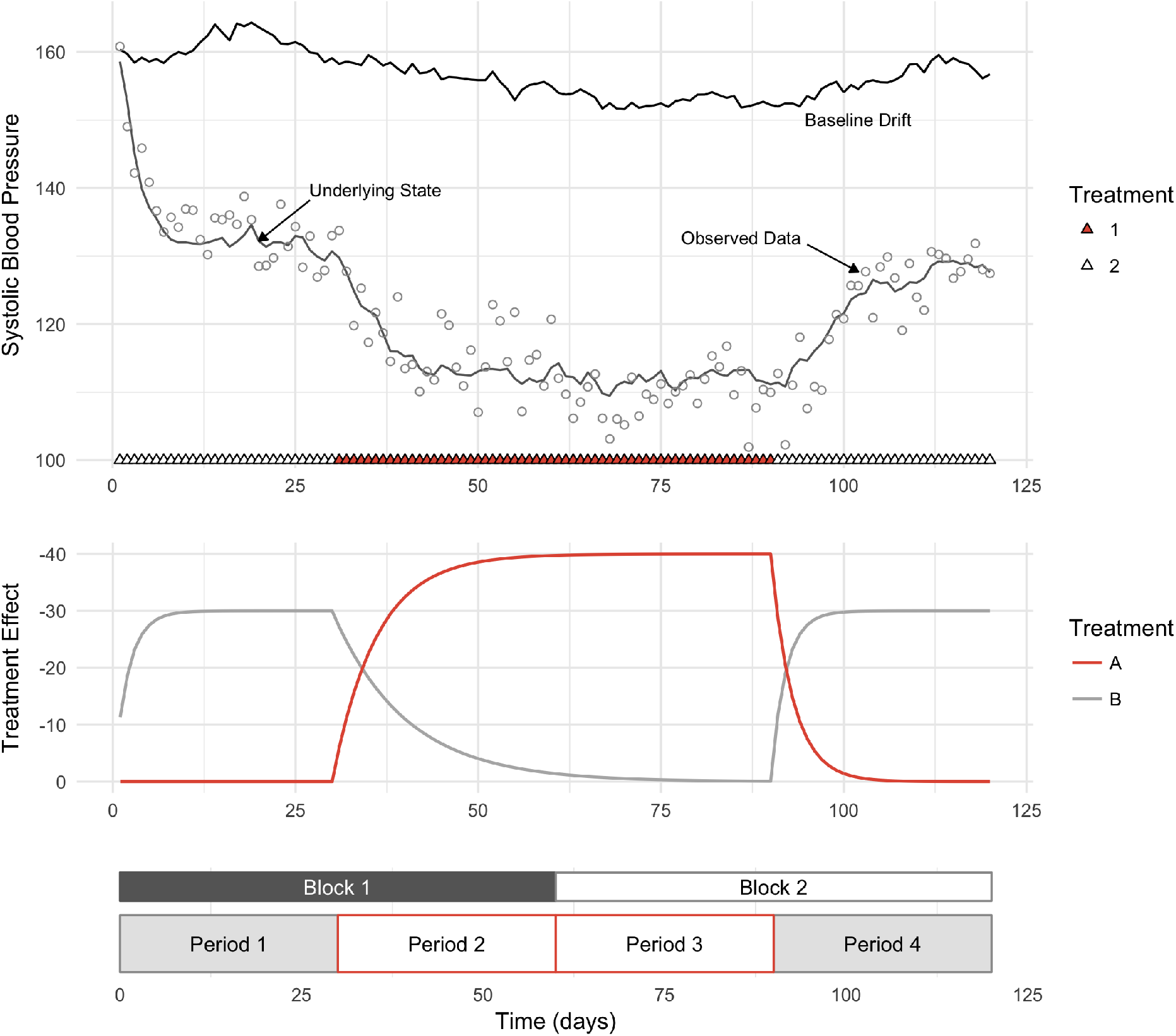
Example of an N-of-1 study comparing two blood pressure medications. An N-of-1 study consists of a set of *N* blocks, each of which contains *J* different treatment periods. The order of the treatment periods within each block is usually randomized. Parameters: *X*_0_ = 160, *E*_1_ = —40, *E*_2_ = –30, *τ*_1_ = 6.0, *γ*_1_ = 3.0, *τ*_2_ = 2.0, *γ*_2_ = 10.0, *α* = 0.5, *P* = 30, *N* = 2, *J* = 2, *σ_b_* = 0.9, *σ_p_* = 1.0, *σ_o_* = 4.0. In this example, one sample was taken per day.

N-of-1 studies inform the care of individual patients while simultaneously generating evidence that can be combined with other N-of-1 studies to yield population-level analyses [9–11]. These studies will likely play a key role in precision medicine, with its focus on narrowly-defined patient cohorts, rare conditions, and complex comorbidities [5].

However, the design and analysis of N-of-1 studies present several methodological challenges. Although the Agency for Healthcare Research and Quality recently released a set of statistical guidelines for N-of-1 studies [6, 8], drawing attention to potential treatment effect confounders like underlying time trends, carryover effects, and autocorrelated measurements, there is currently no universal methodological or statistical framework for the design and analysis of N-of-1 trials. Treatments are often compared graphically, or using ad-hoc measures of efficacy that differ from study to study; a review of N-of-1 trials published between 1985 and 2010 found that only 49% used any statistical measure to compare treatments [2]. As a result, it is difficult to compare findings from different studies or understand how specific analytic choices influence study results.

N-of-1 studies must also overcome daunting practical and logistical challenges. For example, although researchers might like to administer treatments over dozens of blocks to increase statistical power, such designs are burdensome to the patient and increase the likelihood of attrition. It is also difficult to convince individuals to revisit earlier treatments, especially if these are perceived as less effective [1, 6]. Practically speaking, this means the number of treatment blocks in an N-of-1 study is limited, as is the total duration of the study. And although a statistician might prefer more, shorter blocks relative to fewer, longer blocks (since the number of samples in a traditional N-of-1 analysis is linear in the number of blocks), rapid switching among treatments may obscure true differences in efficacy due to carryover effects from earlier treatments. Many treatments, such as antidepressants, also take time to display their full effects. Decisions about the length and arrangement of treatment periods can have a profound effect on statistical effect estimates in N-of-1 studies.

Simulation has played a crucial role in clinical trial design, increasing the efficiency and cost-effectiveness of clinical trials, especially in the pharmaceutical industry [12]. Inspired by this, we have developed a stochastic time series simulation model for N-of-1 studies that incorporates all study-relevant effects, such as carryover and wash-in effects. The model can be used to produce realistic simulated data for a near-infinite number of N-of-1 study designs, treatment profiles, and patient characteristics. The model also incorporates noise parameters like baseline drift, short-term fluctuations (process noise), and measurement error to provide realistic sources of variation that can obscure treatment effect in real patient settings. Using simulation, we can cheaply and easily investigate how design parameters like sampling frequency, number and location of samples within blocks, treatment order within blocks, treatment period duration, and total number of blocks impact statistical estimates of treatment effect.

Here we use the model to analyze two N-of-1 “case studies”, showing how simulation can both optimize study designs and assist researchers in deciding on an appropriate analysis protocol. We then use the model to produce a set of design recommendations for N-of-1 studies based on parameters of the treatment, outcome, and instrument. We provide our simulation software as a supplement to the paper.

## Results

### Modeling the Key Features of an N-of-1 Study

The complete set of parameters for our model can be found in Table 1. The basic model consists of an underlying deterministic process (the growth and decay of treatment effects over time) in addition to three types of noise: random baseline drift (e.g. long-term illness onset and recovery processes, gaining/losing weight, long-term changes in blood pressure, etc.), process noise, which manifests as short-term fluctuations (e.g. heart rate and blood pressure volatility, periods of activity/inactivity, changes in sleep and diet from day to day), and observation noise, which is a function of the instrument and is not related to any underlying biological effect (e.g. the measurement noise associated with the cuff that is used to monitor blood pressure).

**Table 1:** The parameters underlying data generation for an N-of-1 study. The parameters are divided into study design parameters (D), treatment-related parameters (T), measurement parameters (M), and outcome-related parameters (O).

| Parameter | Type | Description |
| --- | --- | --- |
| $\{t_1, \dots, t_n\}$ | D | sampling times |
| $N$ | D | number of blocks (each with $J$ periods in random order) |
| $J$ | D | number of treatment periods per block |
| $P$ | D | treatment period length |
| $E_1, \dots, E_J$ | T | effect sizes for treatments 1 through $J$ |
| $\tau_1, \dots, \tau_J$ | T | run-in time constants for treatments 1 through $J$ |
| $\gamma_1, \dots, \gamma_J$ | T | wash-out time constants for treatments 1 through $J$ |
| $\alpha$ | O | sensitivity to treatment effect |
| $\sigma_b^2$ | O | variance of baseline drift process |
| $\sigma_p^2$ | O | variance of process noise |
| $\sigma_o^2$ | M | variance of observation noise |

We divide the parameters into four groups: *study design parameters,* which the study designer can vary, *treatment parameters,* which are immutable features of the particular treatments under consideration, a *measurement parameter,* which is a feature of the device used to measure the outcome, and *outcome parameters,* which are features of the underlying biological process under consideration and may vary from individual to individual. A diagram of an N-of-1 block design and our model of how treatment effects vary over time is in Figure 1.

### Case Study: Optimizing Study Design

Simulation allows us to investigate the impact of subtle design choices on the likelihood of study success. To illustrate this, we simulated a study of two different blood pressure medications and their impact on systolic blood pressure, similar to the data shown in [5] (see Methods for details). The study parameters, underlying (unobserved) data, and observed data are in Figure 1. The results of several hundred simulations of this study are shown in Figure 2. We used one of the standard N-of-1 regression models outlined in [6] and [8] to estimate treatment effect and obtain an associated *p*-value.

**Figure 2:**
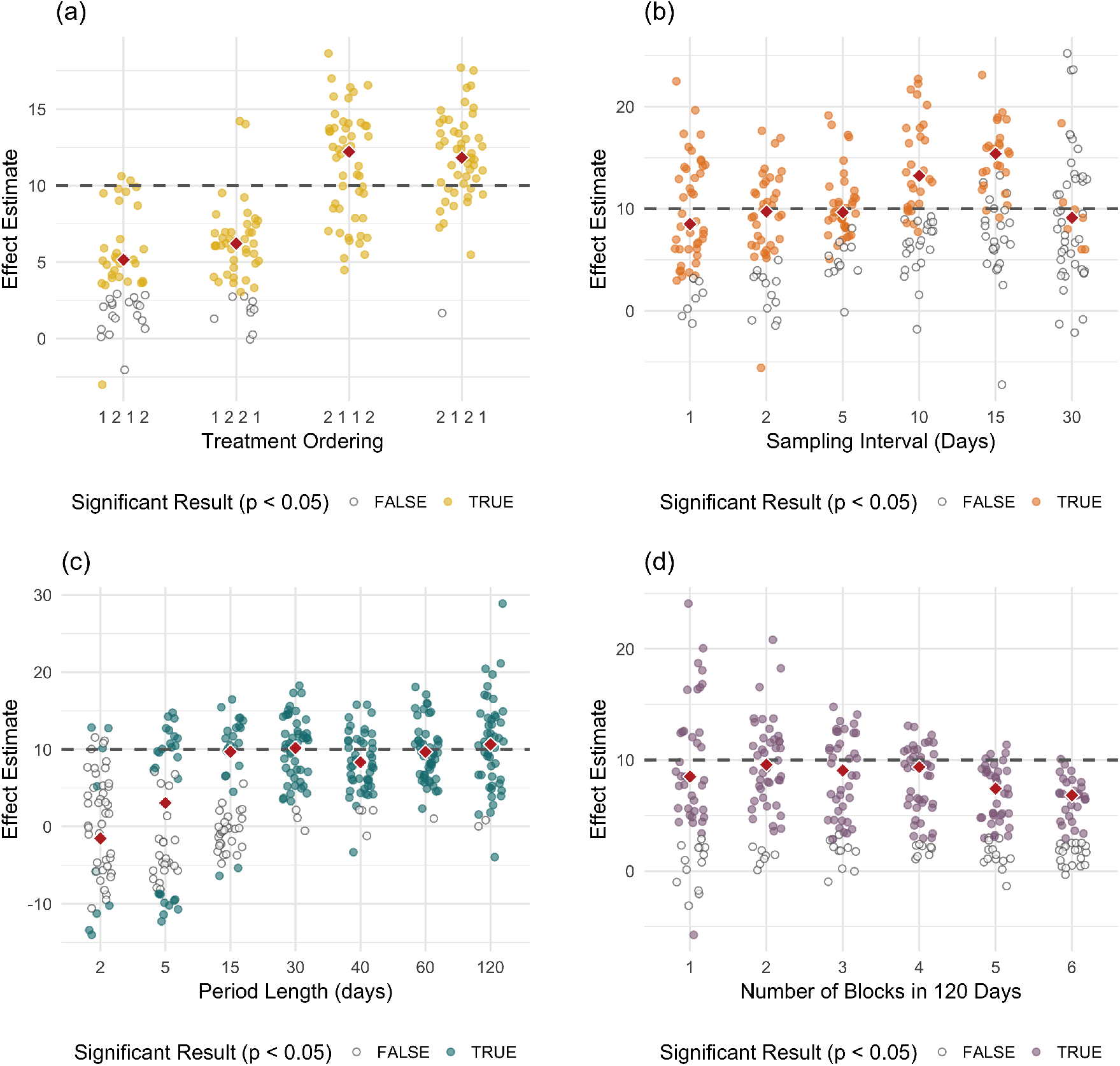
Variation in effect estimates for the hypertension study by study design parameters, including (a) treatment period ordering, (b) sampling frequency, (c) treatment period length, and (d) number of blocks for a fixed study length. The true effect size is 10, illustrated by the dashed lines in the figures. The red diamonds correspond to the median effect size for the statistically significant results within each group. Power estimates were obtained by calculating the ratio of the number of colored dots to the number of total dots. There are 50 trials shown for each parameter setting.

The ordering of treatment periods has a strong effect on both statistical power and effect size estimates (Figure 2a). When treatments are administered in the order *2 12 1* or *2 112,* power – at a standard 5% significance level – is at or near 1.0 (a difference in effects between the two treatments is nearly always detected) and the median effect size estimate is close to the true value of 10.0. In contrast, when the sequence *12 12* is used, power drops to approximately 0.6 and the median difference in effects is 5.8 ± 2.9, about 60% of the true value. This is because Treatment 1 takes longer to reach its full effect than Treatment 2, and the patient starts at a relatively high baseline (systolic bp = 160), so Treatment 1 never reaches its full effect during the first treatment period before the switch to Treatment 2.

Increasing the sampling frequency (Figure 2b) causes power to increase, but only to a point. When only one sample is taken at the end of each treatment period, which is the most common approach to analyzing N-of-1 studies [6, 8], power is only 0.14. Sampling every day, predictably, yields a much increased power of 0.84. However, sampling every 2 or 5 days could substantially reduce patient burden while causing only a modest reduction in power (to ≈ 0.75).

Figure 2c shows the effect of treatment period length, keeping the total number of blocks fixed at 2 and the sampling rate fixed at 1 sample/day. With a period length of 30 days, one obtains as accurate an effect estimate, on average, as a period length of 60 days while shrinking the total study duration from 240 to 120 days. Period lengths that are too long also run the risk of higher variance in estimates due to baseline drift, as we see with a period length of 120 days in Figure 2c.

Finally, Figure 2d shows the effect of different block designs for a study of fixed length (120 days). Using 3 — 4 blocks appears to be the best approach, as this reduces variance in the effect size estimate relative to a single-block study. Adding more than 4 blocks increases the impact of wash-in/carryover effects on the estimate, which deviates further from its true value of 10 with each additional block.

### Case Study: Evaluating Analysis Protocols

Simulation can also help us evaluate the likely success of new analysis protocols and decision criteria for N-of-1 studies. We simulated a previously-published study [13] in which the outcome was a “diary score” on a scale of 0 to 6, with 0 representing “no complaints at all” and 6 “unbearable complaints”. The study design used 5 blocks, each with 2 treatment periods; only data from the last week of each treatment period was analyzed.

In the paper, the data were analyzed as follows: the researchers took differences in median diary scores between NSAID and paracetamol treatment periods in each block, then calculated number of treatment blocks for which the NSAID score was at least one point lower than the paracetamol score for the patient’s main complaint. An NSAID was recommended if this was true in at least 4/5 blocks. We refer to this method as “median differencing” from now on.

We compared median differencing to the same regression model used in the previous section [8]. Simulations show that median differencing is much more conservative in recommending an NSAID than a standard regression model trained on the same data (Figure 3). For a true effect difference of size 2 (NSAID reduces pain by 2 points relative to paracetamol), median differencing will only recommend an NSAID, on average, 61% of the time, compared to 100% of the time for the regression model. In addition, median differencing will recommend an NSAID more frequently in cases where the diary score on paracetamol is already low (the patient is not in much pain); when the score is high, it becomes harder for it to detect an effect. For a patient with a paracetamol diary score of 6 (the maximum possible pain), if the NSAID reduces the diary score to 4, median differencing will only recommend an NSAID 30% of the time, as opposed to 100% of the time for the regression model. The difference between the two models is even more pronounced when the NSAID only reduces the pain score by 1; in that case, median differencing will only recommend an NSAID, on average, 7% of the time, as opposed to 92% of the time for the regression model.

**Figure 3:**
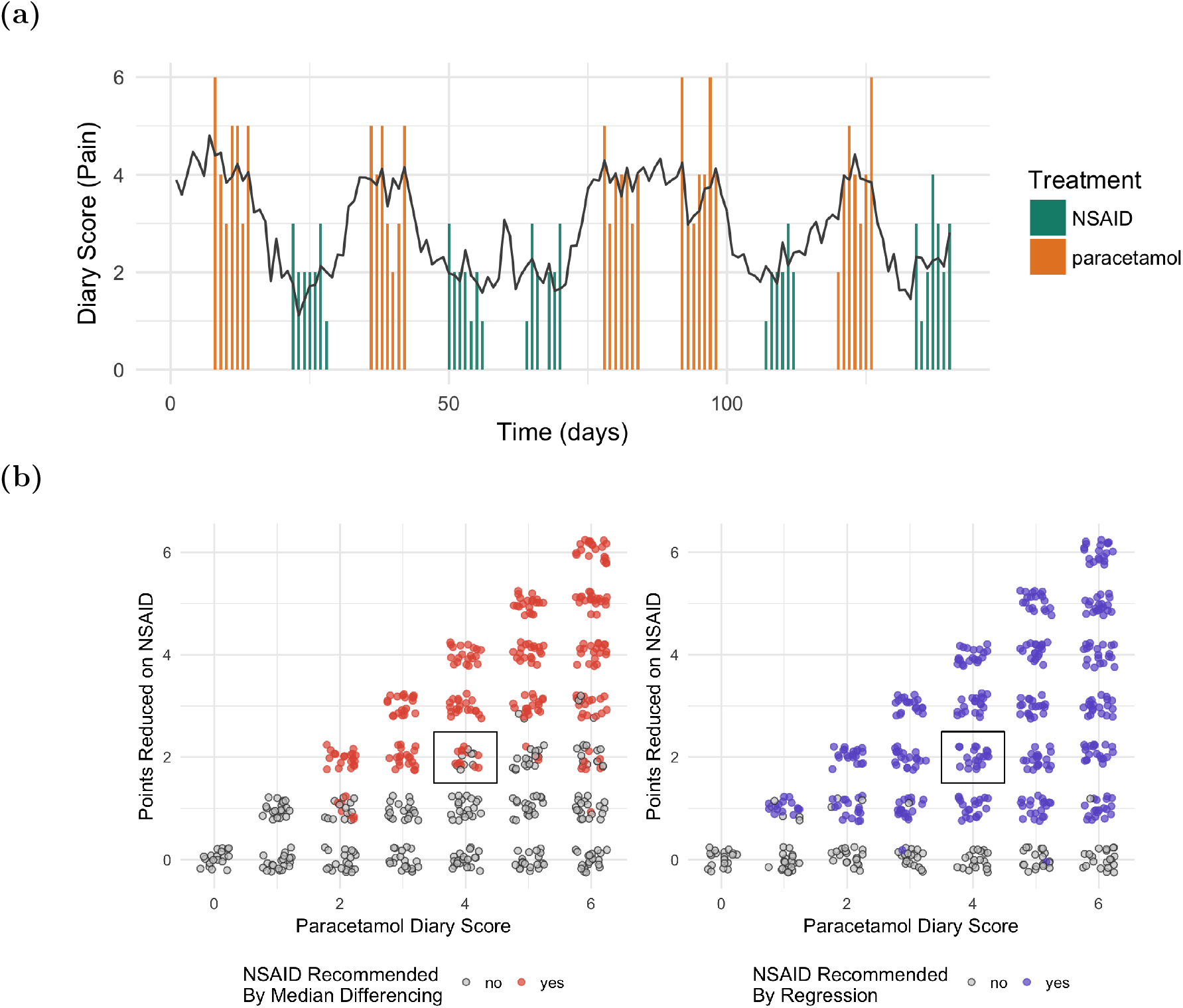
Analyzing a published N-of-1 study comparing NSAIDs to paracetamol. (a) An example simulation in which the true diary score on the NSAID is 2 and on paracetamol is 4. The black line shows the simulated mean outcome (unobserved) at each timepoint, and the colored bars show the observed data, which is a discrete score between 0 and 6. (b) A comparison of the new analysis method with a standard regression model. At the noise levels and effect sizes shown in (a), the new method will recommend an NSAID only about 60% of the time (black rectangle), whereas a regression model will recommend it 100% of the time. Model parameters: *τ*_1_ = *τ*_2_ = 1.0 day, *γ*_1_ = *γ*_2_ = 3.5 days, *α* = 1.0, *σ_b_* = 0.0 (no baseline drift), *σ_p_* = 0.5, *σ_o_* = 1.0.

### Design Considerations for N-of-1 Studies

Figure 4 shows the results of a set of simulations based on “best-case scenarios” – zero variation in parameters other than those under investigation, as well as instantaneous treatment effects (i.e. no carryover effects). The technical details of the simulations can be found in the Methods. Their unifying theme is that they relate study design parameters to (a) statistical power – the ability to detect a treatment effect difference if one exists, and (b) the accuracy of the effect size estimate produced by the model. All compare a single treatment against placebo.

**Figure 4:**
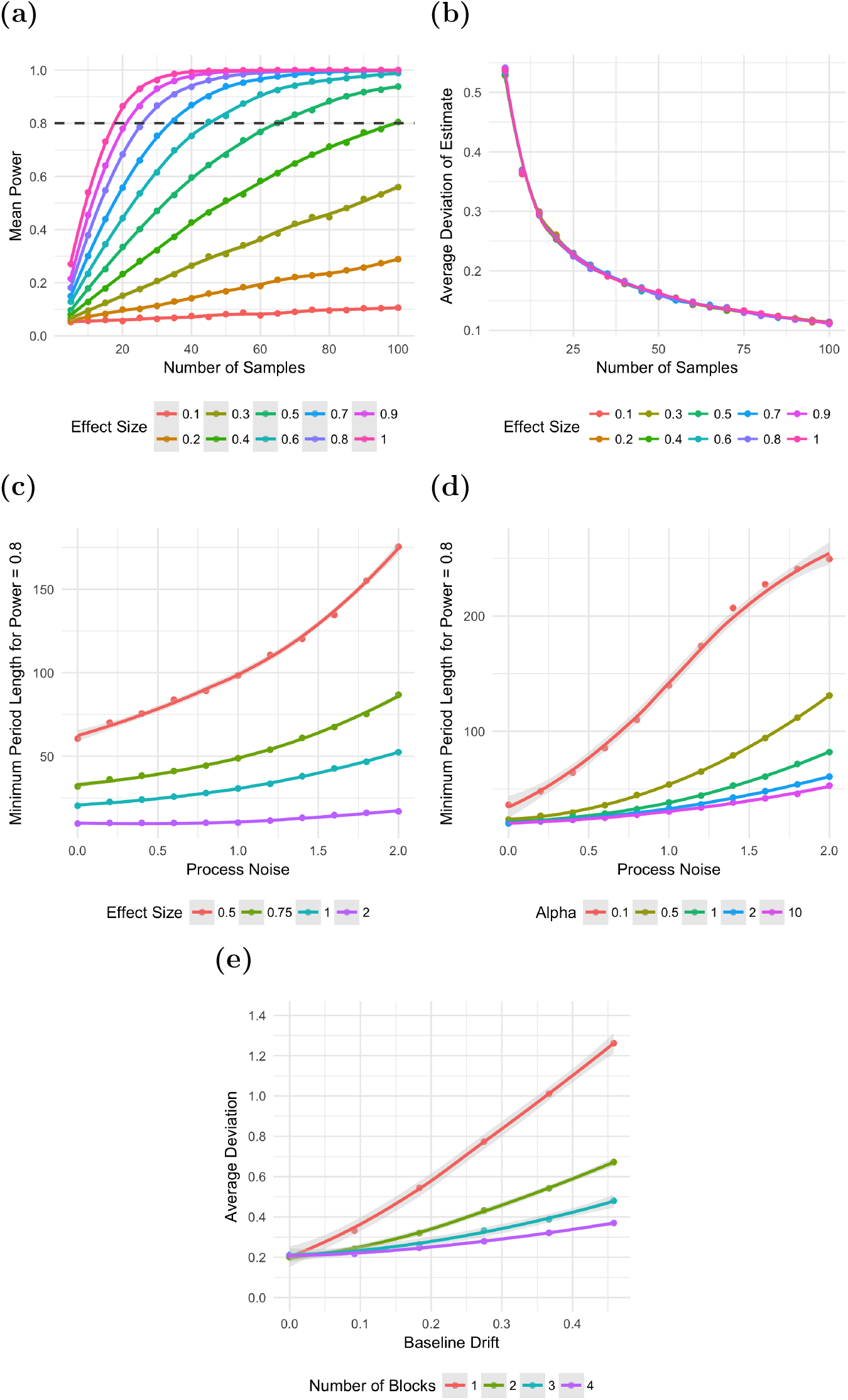
Examining the effect of study design choices on power and accuracy of effect size estimates for an N-of-1 study with effectively instantaneous transitions between treatment states. (a) Effect size vs. power for fixed observation noise (*σ_o_* = 1.0) and no process noise or baseline drift. (b) Average deviation of estimate from true value vs. effect size for fixed observation noise (*σ_o_* = 1.0) and no process noise or baseline drift. (c) Minimum treatment period length (i.e. number of samples, with sampling rate fixed at 1 sample/time unit) required to attain power = 0.8, for varying degrees of process noise and varying effect sizes. No observation noise or baseline drift is present. (d) Same as (c) except effect size is fixed at 1.0 and *α* (individual treatment response) is varied. (e) Average deviation of effect size estimate from its true value, as a function of baseline drift and number of blocks. The effect of baseline drift on the estimate is much more pronounced when fewer blocks are used.

In Figures 4a and 4b, observation noise σ_o_ is fixed at 1.0, with no process noise or baseline drift. As a result, “effect size” really describes a signal-to-noise ratio and is treatment and instrument agnostic. We observe that this ratio impacts power (Figure 4a) but not the accuracy of the effect estimate (Figure 4b).

When the effect size is 1.0, a minimum of about 20 samples from each treatment are needed to conclude that a treatment effect exists (at a standard 5% significance level and power = 0.8). When the effect size drops to 0.5, approximately 65 samples per treatment are required. Even more samples will be needed in real-life situations where process noise, baseline drift, and carryover effects all play a role. This indicates that unless the effect size is very high relative to the observation noise, N-of-1 studies using only a few blocks, with a single sample taken per block (the traditional approach to analyzing N-of-1 studies) will be vastly underpowered.

A separate consideration is the error in the effect size estimate, which declines monotonically with the number of samples (Figure 4b). To obtain an estimate within 0.2*σ_o_* of the true estimate, at least 30 samples per treatment are needed; to reach 0.1*σ_o_*, over 100 samples per treatment are needed.

Figure 4c shows the impact of process noise on the samples needed to attain power ≥ 0.8, in the absence of observation noise and baseline drift. Here the inter-sample interval is fixed at 1 sample/time unit and the process noise is defined relative to that; *σ_p_* = 1.0 means that if no treatment effect were present, the variance of the Wiener process underlying the process noise would be 1 outcome unit/time unit. We see that regardless of effect size, increasing the process noise from 1.0 to 2.0 roughly doubles the number of samples it takes to attain a power of 0.8. However, the effect is nonlinear; below *σ_p_* ≈ 1.0, the number of samples needed flattens out in the absence of other sources of noise.

In Figure 4d, we see the impact of individual sensitivity to treatment effect on study outcome. The lower *α* is, the less effect changes in treatment have on the outcome relative to random fluctuations due to process noise. We see this when we contrast the effect of increased process noise on the minimum samples required under conditions of low (α = 0.1) and high α = 10.0 treatment sensitivity.

Finally, Figure 4e shows us why we bother to have blocks at all: to guard against baseline drift. The figure shows what happens in a study of total length 240 days when block designs incorporating 1, 2, 3, or 4 blocks are used. As baseline drift increases (holding process and observation noise constant at *σ_p_* = *σ_o_* = 0.0), the effect size estimate provided by the model increasingly deviates from its true value. This effect is most pronounced in studies with only a single block, and decreases as the number of blocks increases.

## Discussion

### Recommendations for the Design of N-of-1 Studies

An N-of-1 study should have as many blocks as possible to avoid baseline drift (Figure 4e). In theory, if no wash-in or carryover effects are present, the ideal situation is for the number of samples to be *JN*, where *N* is the number of blocks and *J* the number of treatments, and a single sample should be taken at the end of each treatment period. In practice, however, the number of blocks we can use in a study is bounded by the dangers of administering different treatments in rapid succession, the time it takes treatments to ramp up to their full effects (“run-in”: Table 1), the time it takes them to stop working when they are discontinued (“wash-out”: Table 1), and participant patience.

It is important to consider the fact that most N-of-1 studies of reasonable length and reasonable sampling frequency will be underpowered unless the difference in treatment effects is at least on the order of the standard deviation of the observation noise (Figure 4a). The goal, perhaps obvious, should be to measure the outcome with as little noise as possible and at as high a frequency as possible, and/or to continue the study until enough samples are obtained to ensure that the effect will be detected if it is there.

Finally, it is important to remember the difference between power and accuracy. Just because a statistically significant difference in treatment effects is detected, it doesn’t mean that the *quantitative* estimate of *E*_2_ – *E*_1_ reported by the model is accurate. Even when a study is sufficiently powered, the effect size estimate will always improve with the addition of more samples.

Beyond these general statements, our main recommendation for N-of-1 study designers is: simulate your study. We can see from Figures 4c and 4d that process noise and individual sensitivity to treatment can have a dramatic impact on the number of samples needed to adequately power a study, especially if the effect size is small. Simulation can help avoid costly mistakes and maximize the chance of the patient’s being provided with a useful treatment recommendation (if there is a difference) or at least a confident “it doesn’t matter which one you use”.

On a final note, the importance of simulation also extends to analysis methods. As we see in Figure 3, the somewhat arbitrary nature of how many N-of-1 studies are analyzed has the potential to create different sources of bias. It’s important to compare novel analysis methods to the standard models provided by AHRQ and others [6, 8] so study designers are aware of when each can fail. In the paper we analyzed in Figure 3, the authors commented that three months after the end of their study, four of the six patients for whom paracetamol had been recommended were taking NSAIDS, often because they perceived paracetamol to be insufficient to manage their pain despite the results of the study. Although it’s impossible to know for sure, it could be the case that simply by choosing a different approach to data analysis, the researchers would have recommended NSAIDs more frequently. Simulations can help us evaluate these types of issues in advance.

### Modeling Different Outcome Types

Most of our analyses in this paper concerned a continuous (or near-continuous) random variable, such as blood pressure or heart rate. However, many N-of-1 trials use outcomes that are better modeled as counts, proportions, binary random variables (yes/no), or discrete, bounded scores (such as surveys). We can simulate studies of these types by transforming the output of the stochastic differential equation model using a set of transformations similar to those for generalized linear models (see Table 2). We used one such transformation to discretize the scores for the pain management case study.

**Table 2:**
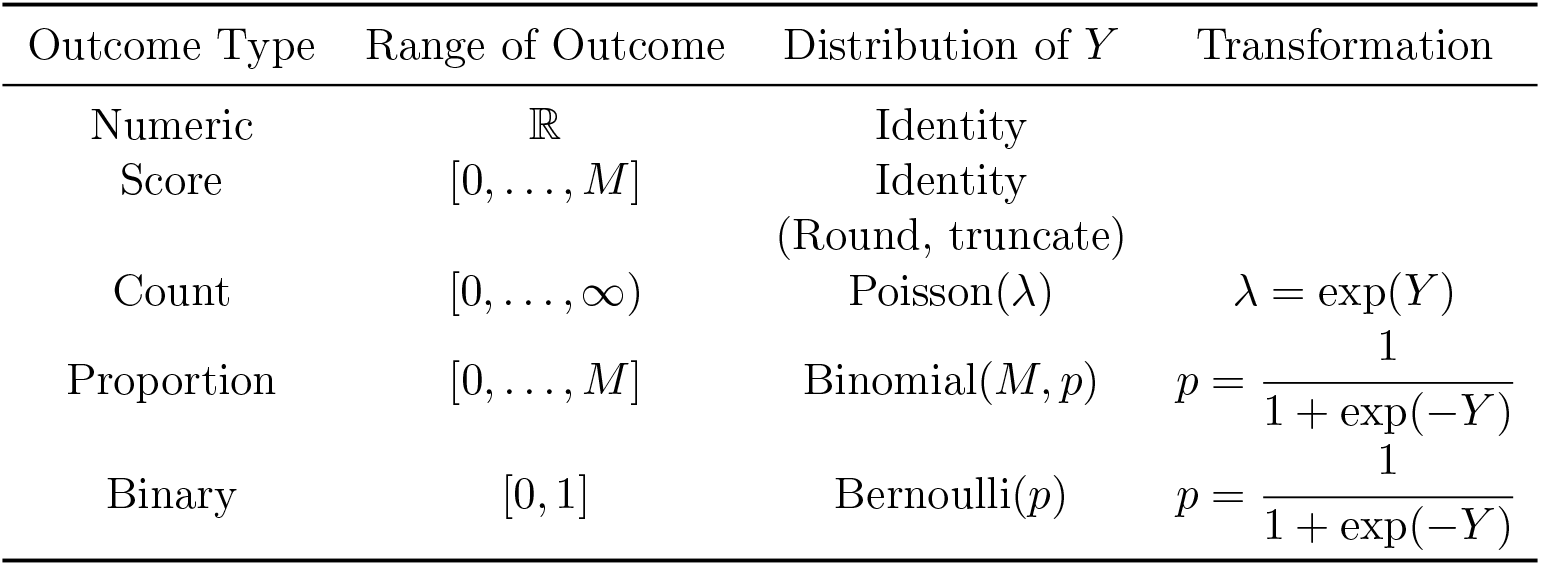
Suggested transformations of *Y* for simulating discrete outcomes.

| Outcome Type | Range of Outcome | Distribution of $Y$ | Transformation |
| --- | --- | --- | --- |
| Numeric | $\mathbb{R}$ | Identity | |
| Score | $[0, \dots, M]$ | Identity<br>(Round, truncate) | |
| Count | $[0, \dots, \infty)$ | Poisson($\lambda$) | $\lambda = \exp(Y)$ |
| Proportion | $[0, \dots, M]$ | Binomial($M, p$) | $p = \frac{1}{1 + \exp(-Y)}$ |
| Binary | $[0, 1]$ | Bernoulli($p$) | $p = \frac{1}{1 + \exp(-Y)}$ |

### Sources of Treatment and Instrument Parameters

By far the strongest drawback to the simulation approach is the difficulty associated with identifying reasonable simulation parameters, especially in cases where the outcome is not a continuous value (see Table 2).

Some parameters have relatively clear interpretations and can be found by looking at known characteristics of treatments and instruments. For example, in the case of a continuous-valued outcome, we can think of the treatment effect, *X_j_*(*t*), as the treatment’s maximum impact – at each point in time – on the outcome in the absence of any noise, in a population of people exactly like the one who is undergoing the study. The treatment effect is governed by three parameters: *τ_j_*, the time constant of “wash-in” for that treatment, *γ_j_*, the time constant of “wash-out”, and *E_j_*, the asymptotic effect size (the change from baseline that the person would experience in the long run, were he/she to continue on this treatment). In the case of a pharmaceutical intervention, these are important parameters that have probably been estimated in earlier clinical trials and used to guide dosages, dosing frequencies, etc. Similarly, reasonable values for *σ_o_* can often be obtained from technical specifications of whatever instrument is used to measure the outcome.

The emerging field of mobile health may provide some help in estimating parameters like *σ_p_* and *σ_b_*, which are properties of an outcome and its natural variation over time [14]. As we begin to monitor patients longitudinally with increasingly higher resolution, our quantitative understanding of long and short-term variation in biological processes will naturally increase. In simulations at present, however, we recommend experimenting with varying parameter scales and examining raw plots of the data to see if the level of noise produced by the model is reasonable. It may also make sense to test ranges of *α, σ_b_*, and *σ_p_* and examine plots like those shown in Figure 4 to assess the effect of these parameter choices on statistical models.

### Study Limitations and Future Work

This study fits simulated data with the simple regression model recommended by the AHRQ, but the data themselves are simulated using a more realistic model. A natural next step would be to use the full simulation model as the basis for fitting data. Future versions of our software will allow users to fit data using the AHRQ model and the full time-series model in a Bayesian framework, which infers parameters using posterior probability distributions given the data rather than point estimates [16, 17]. Thus, uncertainty is an inherent part of the model. This will provide a basis for directly comparing the performance of the full time-series model against the simple AHRQ model for making treatment recommendations. Additionally, posterior parameter distributions inferred from real data can be used to generate more realistic simulated data. This will be especially useful for studies with discrete outcomes, where the linkage between model parameters and outcome data is more difficult to interpret.

Another advantage of a Bayesian parameter estimation approach is that is allows parameter estimates for N-of-1 studies to be continually updated as more individuals undergo the same study, creating a system that learns from past data to adapt the design of future studies.

It is also possible that long-term seasonal, day of week, and time of day effects can influence the outcome of N-of-1 studies. Future versions of our model may incorporate parameters for these effects and fit them using methods akin to those of Prophet [15] or other Bayesian time-series models.

In general, the development of realistic simulations of N-of-1 studies is an ongoing process. We believe that simulation will prove crucial as N-of-1 studies enter mainstream clinical practice, especially in the realm of precision medicine, and we hope that our model will inspire others to adopt N-of-1 studies as a tool in their own research.

## Methods

### Stochastic Time-Series Model

Assume there are *J* total treatments in an N-of-1 study. Let *B*(*t*) denote the patient’s true baseline at time *t*. Let *X_j_*(*t*) denote the effect of treatment *j* (*j* = 1,…, *J*) at time *t*, so that the total treatment effect at time *t* is *X* = *Σ_j_ X_j_* (*t*). Let *T_j_*(*t*) be 1 if treatment *j* is in process at time *t* and 0 otherwise (see Figure 1). Let *Z*(*t*) denote the patient’s true outcome state at time *t*, and let *Y*(*t*) denote the patient’s observed outcome at time *t*.

The underlying effect driver for each treatment is described as an ordinary differential equation:

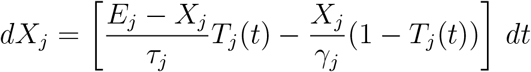

Here each *X_j_*(*t*) is an exponential decay toward a target value that changes over time – either *E_j_* or 0, depending on *T_j_*(*t*) – with time constant *τ_j_* during run-in (decay toward *E_j_*) and *γ_j_* during wash-out (decay toward 0).

Baseline drift is simulated as a discretized Wiener process, where normal noise with variance 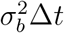 is applied every *Δt*:

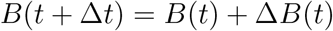

where

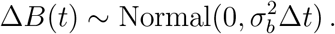

The outcome variable *Z*(*t*) is also a discrete-time stochastic process,

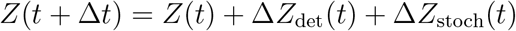

where Δ*Z*_det_(*t*) is a deterministic exponential decay toward the target *X_j_* (*t*) + *B*(*t*):

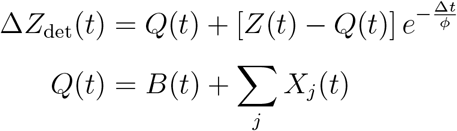

with time constant *φ*, and

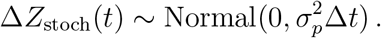

The observed outcome differs from the true outcome only through the addition of normally-distributed observation noise:

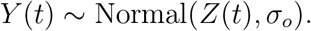

Transformations of *Y*(*t*) can be used to model different types of outcome parameters, such as scores, counts, and binary outcomes (Table 2).

### Hypertension Case Study

A sample dataset and all parameter values for the hypertension case study can be found in Figure 1. The study involves two different blood pressure medications, one of which reduces systolic blood pressure by 10 more points than the other in the long run. The more effective medication, treatment 1, takes longer to reach its full effect (*τ*_1_ = 6.0, *τ*_2_ = 2.0), and less time to wash out (*γ*_1_ = 3.0, *γ*_2_ = 10.0). The sampling rate is 1 sample/day, which we chose to model blood pressure that is monitored using a cuff.

We chose a statistical model for this study that incorporated fixed effects for both block ID and treatment, based on the recommendations provided by the AHRQ and others [6, 8]:

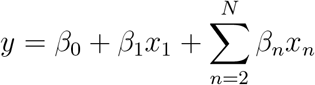

where *x*_1_ is 1 if treatment 2 is in progress at the time of the sample, and 0 otherwise, and *x_n_* is 1 if block *n* is in progress, and 0 otherwise. Note that there are only *n* – 1 indicator variables for blocks; block 1 is used as the reference block.

We experimented with other models, but found that although modeling choices could affect power, effect size estimates did not change much among models. Our software provides the ability to choose from among several different models.

To create Figure 2, we repeated the data generation and analysis process, varying the following parameters and keeping the rest constant:

a. Treatment period orderings were varied among *1 2 1 2, 1 2 2 1, 2 1 1 2,* and *2 1 2 1.*
b. Sampling frequency was varied from 1 sample/day to 1 sample/treatment period, holding the treatment period ordering fixed at *2 1 2 1.*
c. Holding sampling frequency constant at 1 sample/day, period length was varied from 2 to 120 days.
d. Study length was held constant at 120 days and the number of blocks was varied from 1 to 6.

### Pain Management Case Study

The trial design used in this case study was based on the design reported in [13]. Although we did not have access to the raw data for this trial and had to estimate reasonable noise parameters and wash-in/wash-out time constants, our goal was simply to compare the analysis technique from the paper to a more traditional approach involving a regression model with fixed effects for treatment and block [8]. The regression model we chose was the same as for the first case study.

The parameters we chose for this model can be found in Figure 3. We based our decisions about *τ* and *γ* on the fact that the authors chose a wash-out period of one week for the different treatments, and the fact that both NSAIDs and paracetamol are short-acting drugs. We converted the numeric value of the patient state to a discrete score by rounding and truncating it as shown in Table 2.

### Simulations for Design Recommendations

All of the simulations in Figure 4 use a baseline of 0 and time constants (*τ*_1_, *τ*_2_, *γ*_1_, *γ*_2_) of 0.01. Since treatment 1 is assumed to be placebo, its effect size, *E*_1_, is zero. We used *α* = 10 to produce a near-instantaneous effect. Figures 4a and 4b used only a single block, since in the absence of any sources of noise except observation noise, block design doesn’t matter. The rest of the parameter choices are outlined in the figure. Each dot represents an average of 50 trials. The smoothed lines shown in Figure 4 are LOESS fits produced using `geom_smooth` with default parameters in `ggplot`, with spans of 0.4, 0.3, 1.0, 1.0, and 1.0 for subfigures a, b, c, d, and e, respectively.

## Data and Code Availability

The simulation software is available on GitHub at HD2i/n1-simulator. Full details of the available experiments and associated plots are included with the software, along with the datasets generated in the course of making the figures.

## Competing Interests

J.D. holds equity in NuMedii, Ayasdi, and Pillar Health. The rest of the authors declare no competing interests.

## Author Contributions

B.P. and N.Z. jointly conceived of the idea for an N-of-1 simulation model. B.P. and E.B.B. designed the model, wrote the model code, and conducted the experiments for the paper. E.B.B. translated the code into R and created the documentation and user-friendly interface. B.P. drafted the manuscript. M.J., J.D., and N.Z. provided extensive feedback on the manuscript and model design.

## Footnotes

1 or counterbalanced; see [6]

